# A haplotype-resolved chromosome-level genome assembly of *Urochloa decumbens* cv. Basilisk resolves its allopolyploid ancestry and composition

**DOI:** 10.1101/2024.09.25.614935

**Authors:** Camilla Ryan, Fiona Fraser, Naomi Irish, Tom Barker, Vanda Knitlhoffer, Alex Durrant, Gillian Reynolds, Gemy Kaithakottil, David Swarbreck, Jose J. De Vega

## Abstract

Haplotyped-resolved phased assemblies aim to capture the full allelic diversity in heterozygous and polyploid species to enable accurate genetic analyses. However, building non-collapsed references still presents a challenge. Here, we used long-range interaction Hi-C reads (high-throughput chromatin conformation capture) and HiFi PacBio reads to assemble the genome of the apomictic cultivar Basilisks from *Urochloa decumbens* (2n = 4x = 36), an outcrossed tetraploid Paniceae grass widely cropped to feed livestock in the tropics. We identified and removed Hi-C reads between homologous unitigs to facilitate their scaffolding and employed methods for the manual curation of rearrangements and misassemblies. Our final phased assembly included the four haplotypes in 36 chromosomes. We found that 18 chromosomes originated from diploid *U. brizantha* and the other 18 from either *U. ruziziensis* or diploid *U. decumbens.* We also identified a chromosomal translocation between chromosomes 5 and 32, as well as evidence of pairing exclusively within subgenomes, except for a homoeologous exchange in chromosome 21. Our results demonstrate that haplotype-aware assemblies accurately capture the allelic diversity in heterozygous species, making them the preferred option over collapsed-haplotype assemblies.

## Introduction

Livestock contributes to the livelihoods of more than two-thirds of the world’s rural poor (FAOSTAT, 2024). The scarcity and seasonal availability of forage and its low nutritional value are the main limiting constraints in meat and milk production in tropical regions (Bonilla-Cedrez et al. 2023). The use of improved forage cultivars can contribute to building resilience in pasture-based food systems, while boosting animal welfare and the income of rural families. In the broader context, livestock has a large land and carbon footprint, and more nutritious grass can result in lower methane and nitrous oxide emissions per weight (Ferreira et al. 2021; Bonilla-Cedrez et al. 2023). Intensification can also reduce unnecessary degradation of natural land and ecosystem services (Jank et al. 2014).

Native to Africa, *Urochloa* species were introduced into South America in the 1970s because of their good carrying capacity, nutritional value, grazing tolerance, and adaptability to areas of low fertility (Miles et al. 2004; Ferreira et al. 2021). Their broad usage in South America and their ability to interspecifically hybridise have enabled the development of high-performing *Urochloa* cultivars using recurrent selection breeding programmes at CIAT and EMBRAPA (Pizarro et al. 2013; Maass et al. 2015). Improved *Urochloa* varieties have been estimated to double the number of livestock units per area and year compared to natural pastures (Miles et al. 2004; Jank et al. 2014).

Further advances in producing new cultivars are hindered by the genera’s complex genomic architecture, such as variable ploidy, complex phylogenies, apomixis, and a lack of genomic resources (Higgins et al. 2022; Tomaszewska et al. 2023). There are no assemblies for any polyploid *Urochloa* species despite their agronomic importance and the benefits this could bring to breeding programmes (Ferreira et al. 2021). Only single-haplotype genome assemblies were available for the diploid species *U. ruziziensis* (2n = 18) (Worthington et al. 2021), which has limited agronomic interest, prior to this study. These collapsed assemblies are likely not an accurate representation of the species’ haplotypic diversity due to the heterozygous outcrossed nature of the genus.

Until recently, most genome assemblies were collapsed into single haplotypes because algorithms could not distinguish between allelic haplotypes when assembling short-read sequences (Whibley et al. 2021). However, recent advances in sequencing technologies, namely highly accurate long reads and long-range interaction information from chromosomal conformation capture (CCC, Hi-C) methods, are reducing the complexity of genome assembly and enabling the production of haplotype-aware assemblies, including heterozygous and polyploid species (Li and Durbin 2024).

Haplotype-aware, chromosome-level assemblies can greatly benefit crop breeding programmes by enabling more accurate population structure and marker-trait association studies, and a better understanding of gene dosage effects and allelic-driven complex traits, such as apomixis and self-incompatibility (Njaci et al. 2023).

This study presents the first haplotype-aware chromosome-level assembly of the polyploid *U. decumbens* (2n = 4x = 36), particularly from a widely used apomictic cultivar named Basilisk. It demonstrates the feasibility and value of producing fully haplotype-resolved assemblies in heterozygous tetraploid species. We also extended our genome analysis by identifying structural features of *U. decumbens* and clarifying the species’ much-discussed ancestry (Higgins et al. 2022; Masters et al. 2023; Tomaszewska et al. 2023).

## Methodology

### Sample collection

*U. decumbens* cv. Basilisk plants were grown for 8 weeks before DNA extraction from seeds originating from Uganda accessed via the Australian Pastures Genebank [APG 59378] (also known as CIAT 606). After 8 weeks leaf material was harvested following two days in dark conditions. The same sample was used for all the work described below.

### DNA extraction

High molecular weight (HMW) DNA extraction was performed using the Nucleon PhytoPure kit, with a slightly modified version of the recommended protocol. 1g of leaf material was ground under liquid nitrogen for a total grinding time of 9-10 minutes. Following this, the powder was thoroughly resuspended (more aggressively than indicated by the manufacturer protocol) using a 10mm bacterial spreader loop. This method of homogenate mixing was used for all subsequent mixing steps before the addition of the chloroform and resin. After the ice incubation, 300µL of resin was added along with the chloroform. 300µL is at the upper end of the recommended range. The chloroform extraction was followed by extraction with 25:24:1 phenol:chloroform:isoamyl alcohol, which was added to the previous upper phase, mixed at 4°C on a 3D platform rocker for 10 minutes and then centrifuged at 3000G for 10 minutes. The upper phase from this procedure was then transferred to a 15mL Falcon tube and precipitated as recommended by the manufacturer’s protocol. The final elution was left open in a fume hood for two hours to allow residual phenol and ethanol to evaporate, and the DNA sample was left at room temperature overnight.

### Generating HiFi reads

The library for this project was constructed at the Earlham Institute, Norwich, UK, using the SMRTbell® Express Template Prep Kit 2.0 (PacBio®, P/N 100-983-900). 12.6 µg of sample was manually sheared with the Megaruptor 3 instrument (Diagenode, P/N B06010003). The sample underwent AMPure® PB bead (PacBio®, P/N 100-265-900) purification and concentration before undergoing library preparation using the SMRTbell® Express Template Prep Kit 2.0 (PacBio®, P/N 100-983-900). The HiFi library was prepared according to the HiFi protocol version 03 (PacBio®, P/N 101-853-100) and the final library was size fractionated using the SageELF® system (Sage Science®, P/N ELF0001), 0.75% cassette (Sage Science®, P/N ELD7510). The library was quantified by fluorescence (Invitrogen Qubit™ 3.0, P/N Q33216) and the size of fractions was estimated from a smear analysis performed on the FEMTO Pulse® System (Agilent, P/N M5330AA). The loading calculations for sequencing were completed using the PacBio® SMRT®Link Binding Calculator v10.2.

Sequencing primer v5 was annealed to the adapter sequence of the HiFi Library. The library was bound to the sequencing polymerase with the Sequel® II Binding Kit v2.2 (PacBio®, P/N 102-089-000). Calculations for primer and polymerase binding ratios were kept at default values for the library type. Sequel® II DNA internal control 1.0 was spiked into the library at the standard concentration prior to sequencing. The sequencing chemistry used was Sequel® II Sequencing Plate 2.0 (PacBio®, P/N 101-820-200) and the Instrument Control Software v10.1.0.125432. The library was sequenced on three Sequel II SMRT®cell 8M. The parameters for sequencing per SMRT cell were: Adaptive loading default settings, 30-hour movie, 2-hour pre-extension time, 80 pM on plate loading concentration.

### Generating Hi-C reads

Sample material for the Omni-C library prep was 100mg of *U. decumbens* young leaf tissue that was harvested, snap-frozen in liquid nitrogen and stored at −80°C. The Omni-C library was prepared using the Dovetail® Omni-C® Kit (SKU: 21005) according to the manufacturer’s protocol for “Non-mammal v1.2B”. Briefly, the chromatin was fixed with disuccinimidyl glutarate (DSG) and formaldehyde in the nucleus. The cross-linked chromatin was then digested in situ with DNase I (0.05µl). Following digestion, the cells were lysed with SDS to extract the chromatin fragments, which were bound to Chromatin Capture Beads.

Next, the chromatin ends were repaired and ligated to a biotinylated bridge adapter followed by proximity ligation of adapter-containing ends. After proximity ligation, the crosslinks were reversed, the associated proteins were degraded, and the DNA was purified then converted into a sequencing library (NEBNext® Ultra™ II DNA Library Prep Kit for Illumina® (E7645)) using Illumina-compatible adaptors (NEBNext® Multiplex Oligos for Illumina® (Index Primers Set 1) (E7335)). Biotin-containing fragments were isolated using streptavidin beads prior to PCR amplification.

The library pool was diluted to 0.5 nM using EB (10mM Tris pH8.0) in a volume of 18ul before spiking in 1% Illumina phiX Control v3. This was denatured by adding 4µl 0.2N NaOH and incubating at room temperature for 8 mins, after which it was neutralised by adding 5µl 400mM tris pH 8.0. A master mix of EPX1, EPX2, and EPX3 from Illumina’s Xp 2-lane kit v1.5 (20043130, Illumina) was made and 63µl added to the denatured pool leaving 90µl at a concentration of 100 pM. This was loaded onto a NovaSeq SP flow cell using the NovaSeq Xp Flow Cell Dock. The flow cell was then loaded onto the NovaSeq 6000 along with an NovaSeq 6000 SP cluster cartridge, buffer cartridge, and 300 cycle SBS cartridge (20028400, Illumina). The NovaSeq had NVCS v1.7.5 and RTA v3.4.4 and was set up to sequence 150bp PE reads. The data was demultiplexed and converted to fastq format using Illumina Bcl2Fastq2.

### Genome assembly

The workflow used to produce the genome assembly is represented in figure 1. Firstly, a unitig assembly was produced using HiFiasm v0.18 (Cheng et al. 2021; 2022). HiFiasm produces multiple assemblies with increasing contiguity by iteratively improving the assembly graph and increasingly discarding (or collapsing) minor variations. We decided to advance with the unitig assembly (instead of contigs) for scaffolding because unitigs are haplotype-specific; therefore, the unitig assembly included all four haplotypes. The quality of the unitig assembly was comparable to that of the contig level assembly, and the number of unitigs assembled was within the scaffolder’s processing limits (Zhou et al. 2022).

**Figure 1:**
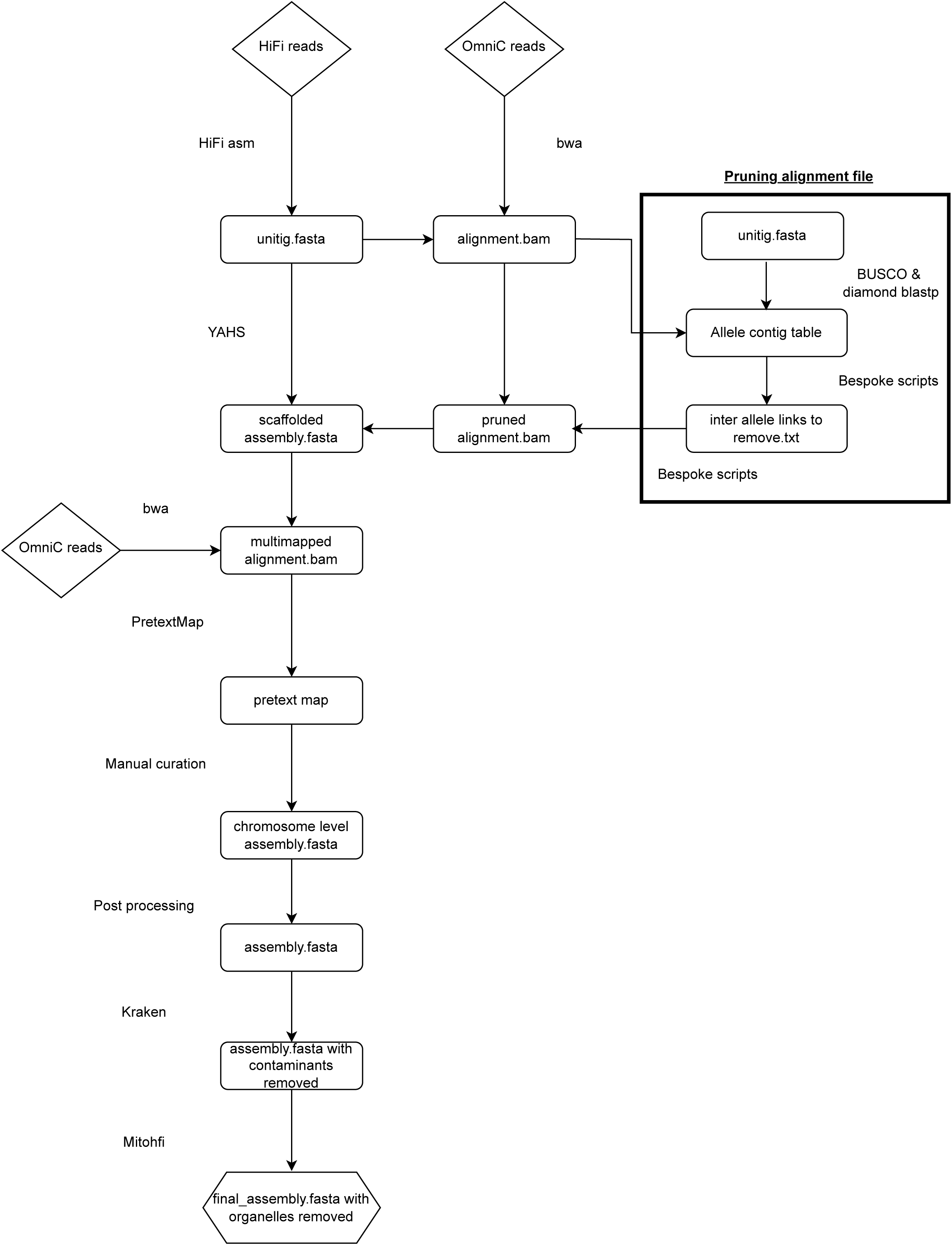
Overview of the bioinformatics pipeline used to create the haplotype-resolved chromosome-level assembly of the allotetraploid *Urochloa decumbens*.

OmniC reads were mapped to the unitig assembly and the read alignment file was pruned for use in scaffolding. Pruning was first suggested by Zhang *et al*. (2018; 2019) and referred to the method of removing uninformative inter-chromosomal links between allelic haplotypes, which otherwise may result in misjoins during scaffolding. For that, the single-copy markers for the grasses family from BUSCO (Manni et al. 2021) were mapped with Diamond blastp (Buchfink et al. 2021) to the unitigs to generate a table of unitigs with the same single-copy marker (named allelic unitigs). Then, the OmniC reads linking allelic unitigs were removed.

The pruned OmniC alignment file was used with YAHS v1.2a.2 (Zhou et al. 2022) to scaffold the unitig assembly with a minimum mapping quality of 30 and the following lists of resolutions 1000, 2000, 5000, 10000, 20000, 50000, 100000, 200000, 500000, 1000000, 2000000, 5000000, 10000000, 20000000, 50000000, 100000000, 200000000, 500000000.

The scaffolded assembly was manually curated following a workflow used by the Darwin Tree of Life programme developed by the Sanger Institute (Cambridge, UK). Firstly, the OmniC reads were remapped to the scaffolded assembly allowing for multi-mapping reads; although multi-mapping reads are not informative during scaffolding, they are useful during manual curation, e.g. to identify regions with repeat sequences. PretextMap v0.1.9 (Tree of Life Programme, 2024a) was used to create a contact matrix (or pretext map) compatible with PretextView v0.2.5 (Tree of Life Programme, 2024b), which is an interactive viewer that allows the editing of the contact matrix directly. Tracks for telomere sequences, gaps and coverage were added to the pretext map to help with manual curation. The location of telomere sequences in the assembly was identified using tidk v0.2.31 (Brown et al. 2023); as the telomere sequences *for U. decumbens* are unknown, the motif “TTTAGGG” was used (Peska and Garcia 2020). Seqtk cutN v1.2 (Li, 2024) was used to localise gaps in the scaffolded genome over 10 bps. Finally, HiFi reads were mapped back to the scaffolded genome using Minimap2 v2.24 (Li 2018; Li 2021) to generate the coverage tracks. The alignment file was sorted with Samtools v1.10 (Danecek et al. 2021) and used to calculate coverage across the genome using Mosdepth v0.3.3 (Pedersen and Quinlan 2018). Tracks were then created from bedgraphs and added to the pretext map using PretextGraph v0.0.6 (Tree of Life Programme, 2024c). After manual curation, the script rapid_pretext2tpf_XL.py (Tracey and Wood, 2024) was used to create a new tpf file and regenerate the genome’s Fasta file. Chromosome-length sequences were sorted and renamed based on homology to each other and subgenome ancestry, and scaffolds by length.

### Contaminant and organelle sequence removal

Kraken2 v2.0.7 (Wood et al. 2019) and the “Standard-16 nucleotide database” version 2.0.7_refseq-201910 (Langmead, 2024) were used to identify any scaffolds that did not belong to Viridiplantae. Mitohifi v3.0.0 (Uliano-Silva et al. 2023) was used to identify scaffolds belonging to mitochondria (mtdna) and chloroplast (pltd) and the most similar sequences available to assemble the two organelles. For *U. decumbens,* the most similar mtdna found was from *Microstegium vimineum* (NC_072666.1) [accessed May 24, 2023]) and the most similar pltd came from *U. decumbens* (NC_030066.1) [accessed May 24, 2023]. Scaffolds identified as non-Viridiplantae (contaminants), chloroplast or mitochondrial were removed from the final assembly.

### Genome quality assessment

The quality of the final assembly was assessed using several measures; basic assembly metrics, including contiguity, were produced using Abyss v1.9.0 (Simpson et al. 2009), assembly completeness was assessed from BUSCO v5.3.2 (Manni et al. 2021) analysis using the Poales v10 database. Merqury v1.3 (Rhie et al. 2020) was used to produce Kmer completeness metrics and Kmer spectra plots to ensure the assembly captured all the content from the reads and to produce consensus quality (QV) metrics. QV measures likely assembly errors based on Kmers found only in the reads. This is then converted into a Phred-equivalent score (Rhie et al. 2020). The final chromosomes were mapped, using Minimap2 v2.24 (Li 2018; Li 2021), to an existing reference of the closely related diploid *U. ruziziensis* [GCA_015476505.1, accessed March, 2023].

### Gene annotation

Gene models were generated from the *U. decumbens* assembly using *Robust and Extendable eukaryotic Annotation Toolkit* (REAT) v0.6.1 (REAT, 2024) and Minos v1.8.0 (MINOS, 2024), which are pipelines that used Mikado v2.3.4 (Venturini et al. 2018), Portcullis v1.2.4 (Mapleson et al, 2018) and multiple third-party tools (listed in the above repositories) as dependencies. Identification of repetitive elements was performed using the EI-Repeat pipeline v1.3.4 (EI-Repeat, 2024), which masked the genome assembly using RepeatMasker v4.0.7 (Tarailo-Graovac and Chen 2009) and the RepBase database and a de novo repeat database constructed with RepeatModeler v1.0.11 (Smit and Hubley 2008). REAT’s “transcriptomic workflow” was used for alignment of short-read RNA-Seq data generated in a previous study (Higgins et al. 2022) using HISAT2 v2.2.1 (Kim et al. 2019) with high-confidence splice junctions identified using Portcullis v.1.2.4 (Mapleson et al. 2018). Alignments from short-reads were assembled using StringTie v2.1.5 (Kovaka et al. 2019) and Scallop v0.10.5 (Shao and Kingsford 2017). A consolidated set of transcriptome-derived gene models was generated using Mikado v2.3.3 (Venturini et al. 2018). REAT’s “Homology workflow” was used to align protein sequences from 7 related species (table S1) against the *U. decumbens* assembly. Proteins were aligned using Spaln v2.4.7 (Gotoh 2008) and filtered to remove misaligned proteins. The same proteins were also aligned using miniprot v0.3 (Li 2023) and filtered. The aligned proteins from both alignment methods were clustered into loci and a consolidated set of gene models derived with Mikado v2.3.4.

REAT’s “Prediction workflow” was used to generate a set of evidence-guided gene predictions by training Augustus (Stanke and Morgenstern 2005) with high-confidence gene models from the previous workflows. Four alternative Augustus runs were performed with varying weightings of evidence, which were provided to EVidenceModeler (Haas et al. 2008) along with the transcriptome and protein evidence to generate consensus gene structures.

Genes were also predicted using Helixer (Holst et al. 2023), a deep neural networks (DNN) approach, using its publicly available plant model. The final set of gene models was selected using Minos from the outputs from REAT’s Homology, Transcriptome and Prediction workflows, plus Helixer’s gene models. Gene models were classified as coding, non-coding, or transposable, and with a high or low confidence score, based on the support from RNA-Seq or protein evidence (from the 7 related species plus UniProt’s Magnoliopsida proteins) with previously defined criteria (Grewal et al. 2024).

### Ancestry analysis

Sourmash v4.8.5 (Irber et al. 2024) was used to generate Kmer signatures for all *U. decumbens* and *U. ruziziensis* chromosomes, perform pairwise comparisons between genomes, plot dendrograms and heatmaps. Kmer composition and frequency signatures, comparisons and plots were generated for Kmer sizes 3-21 in increments of 2 and 21-161 in increments of 10. Subgenomic clustering was determined using the “strict cut” criteria in Reynolds et al. (2024). In short, subgenomic clusters are deemed correct if, and only if, all chromosomes belonging to a subgenome are within the same cluster. In the plots, all chromosome numbers are annotated with their subgenome ancestry. *U. ruziziensis* chromosomes [GCA_015476505.1, accessed March 2023] were also annotated with introgression information and number.

Reads from three *Urochloa* species were aligned to the final assembly to clarify its genome composition and ancestry. Reads from the diploid *U. decumbens* were downloaded from NCBI’s sequence read archive [SRR16327313, accessed on May 22, 2023]. Reads from the diploid *U. brizantha* were kindly shared by EMBRAPA (M. Pessoa, *per. Comm*.). Reads from *U. ruziziensis* were downloaded from PRJNA437375. Each set of reads was mapped using Minimap2 v2.24 (Li 2018) and the coverage was plotted in R v3.6.0 (R Core Team 2021) using a modified version of the function plot_coverage() from PafR v0.0.2 (Winter 2020).

### Assessing repeat content

Transposable elements (TE) were identified from the genome *de novo* using EDTA v2.1.0 (Ou et al. 2019). Long terminal repeats (LTR) were extracted from the output of EDTA and the distribution and density of intact LTRs were plotted across the genome in R v3.6.0 (R Core Team 2021).

### Identifying structural changes through synteny

Structural changes were initially identified using the coverage plots and later in greater detail using a syntenic approach. For that, high-confidence protein-coding genes were selected from the annotation results. However, only those coding genes found on chromosomes (*i.e.* not on scaffolds) were retained. A table of all-vs-all protein alignments was created using DIAMOND blastp v2.0.15 (Buchfink et al. 2021). This table of homologous genes and the genome annotation was used with MCScanx v2 (Wang 2022) to identify putative syntenic chromosomic regions and produce a collinearity file. The results of MCScanX were plotted using SynVisio (Bandi and Gutwin 2020).

## Results and Discussion

### Capturing the full allelic diversity in a heterozygous and polyploid grass species

A haplotype-resolved chromosome-level de novo assembly from *U. decumbens* cultivar Basilisk was generated using a combination of HiFi reads and OmniC data. Starting from 7,122 unitig sequences generated by the assembler, 85.9% of the assembly was later successfully anchored into 36 chromosomes and 7,086 unlocalised scaffolds (Table 1). The 36 chromosomes contained 99.2% complete BUSCO markers. A Kmer spectra of the HiFi reads versus the assembly evidence the assembly accurately reflected the raw read content (Figure S1).

**Table 1:** Quality and contiguity metrics for the genome assembly of *U. decumbens*.

| **Statistics** | **Complete genome (n = 7122)** | **Chromosomes only (n = 36)** |
| --- | --- | --- |
| **Total assembly size (Gb)** | 2.879 | 2.474 |
| **N50 contig length (Mb)** | 66.6 | 69.83 |
| **Max contig length (Mb)** | 104.3 | 104.3 |
| **Complete BUSCOs** | 4,860 (99.2%) | 4,859 (99.2%) |
| **Complete and single BUSCOs** | 36 (0.7%) | 59 (1.2%) |
| **Complete and duplicated BUSCOs** | 4,824 (98.5%) | 4,800 (98.0%) |
| **Fragmented BUSCOs** | 5 (0.1%) | 5 (0.1%) |
| **Missing BUSCOs** | 31 (0.7%) | 32 (0.7%) |
| **Merqury QV** | 67.326 | 71.832 |
| **Merqury completeness** | 97.859 | 95.689 |

Chromosomes were numbered according to the contact matrix after manual curation, which allowed us to identify pairs of chromosomes organised into sub-genomes (Figure 2). The distinctive “chain” pattern of contacts between homologous chromosomes within a subgenome and a faint signal between homoeologous pairs indicated an allotetraploid composition, *i.e.* preferential pairing restricted within subgenomes and no evidence of homoeologous exchanges in the contact matrix (Figure 2).

**Figure 2:**
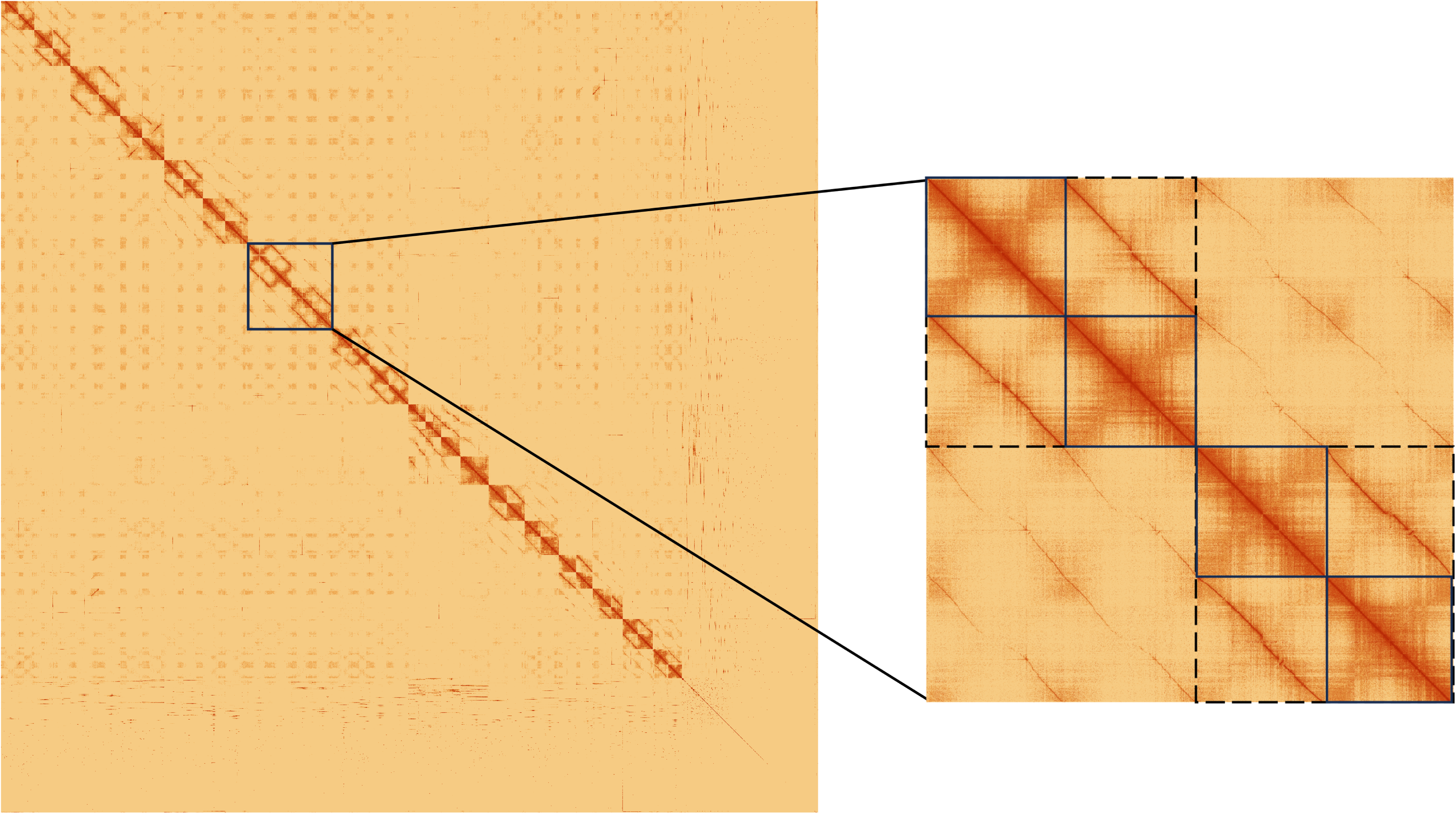
Hi-C contact matrix following manual curation. A close-up of chromosomes 13-16 shows the visual differences between subgenomes (dashed lines) and individual chromosomes (solid lines) within and among subgenomes.

When we aligned the genome to the single-haploid assembly of *U. ruziziensis,* we observed all four *U. decumbens* haplotypes aligned to each *U. ruziziensis* haplotype (Figure 3). For each chromosome, two of the four haplotypes had a lower identity to *U. ruziziensis* than the other two, indicating these chromosomes likely derived from *U. ruziziensis* or a closely related ancestor to it. We also observed evidence of translocations in the dotplot.

**Figure 3:**
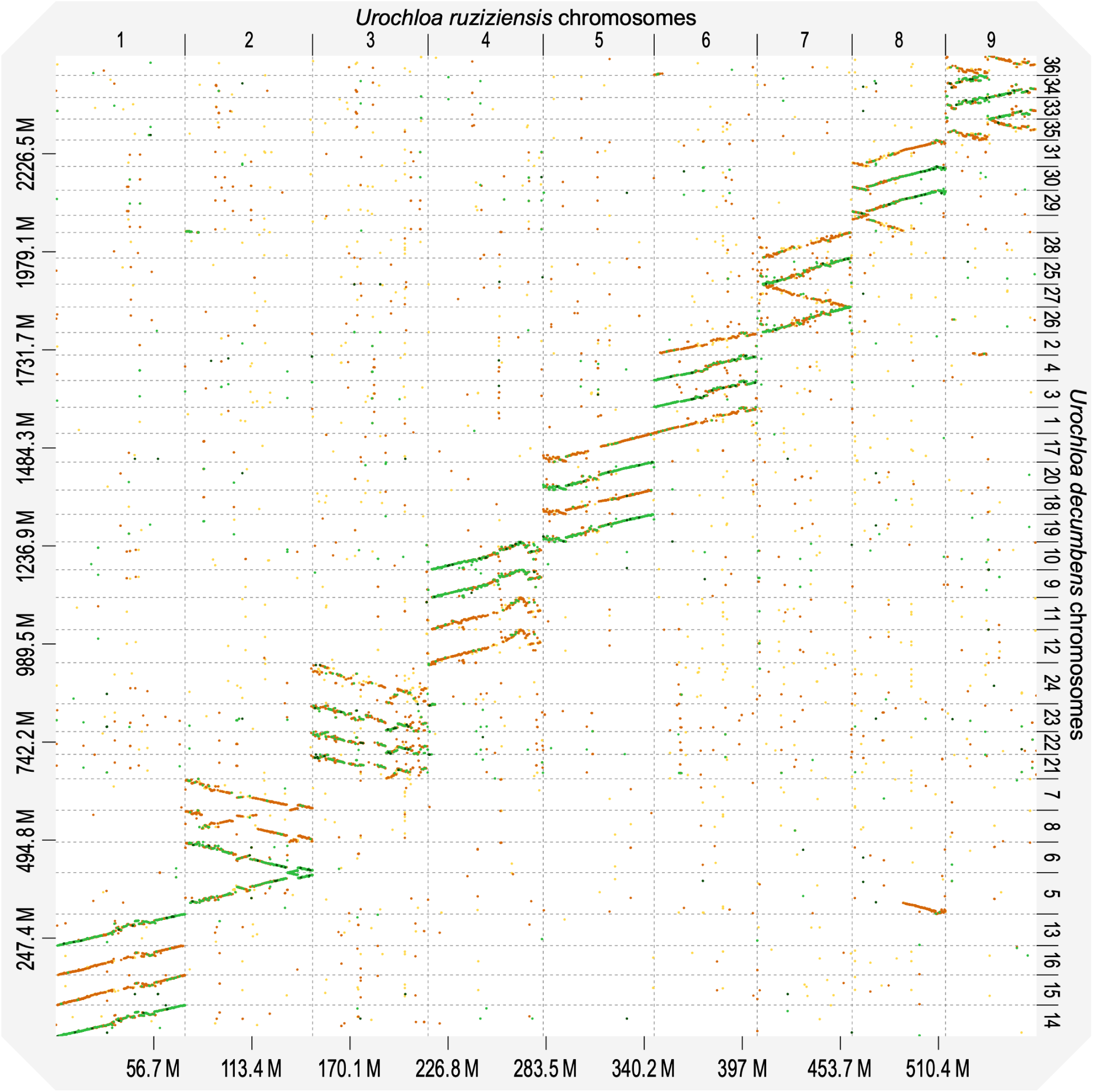
Dotplot representation of the alignments from the 36 chromosomes from allotetraploid *U. decumbens* (4n = 4x =36) to the 9 chromosomes from the single-haplotype assembly of diploid U. ruziziensis (2n = 18). Sequence colours represent alignment identity: 0.00 - 0.25, yellow; 0.25 – 0.50, orange; 0.50 – 0.75, light green; 0.75 – 1.00, dark green.

Out of a total of 4,896 BUSCO markers for Poales, 99.2% were found complete, and 98.5% were duplicated. Similar values were obtained only considering markers found within chromosomes. This indicated that most genic content was captured in the anchored chromosomes. The frequency of alignments showed that most markers aligned four times – which is expected in a complete tetraploid assembly (Figure 7). The REAT annotation pipeline predicted 153,344 protein-coding genes and 207,616 transcripts, with a mean CDS length of 1,732 bp.

### Ancestry of tetraploid *Urochloa decumbens*

*U. decumbens’* chromosomes clustered in two groups by subgenome ancestry (Figure 4a) based on Kmer frequency for most of the sampled Kmer sizes (k=17,21-161; Table S2). The genomes did not cluster by subgenomes based on Kmer composition (Figure 4b) of any k-mer size. Instead, *U. decumbens’* dendrograms reflected the prevalence of preferential pairing within subgenome, except chr. 21 (Figure 4b). I.e. chromosomes clustered in pairs between homologous chromosomes (e.g. Chr13_B, Chr14_B) within the same subgenome.

**Figure 4:**
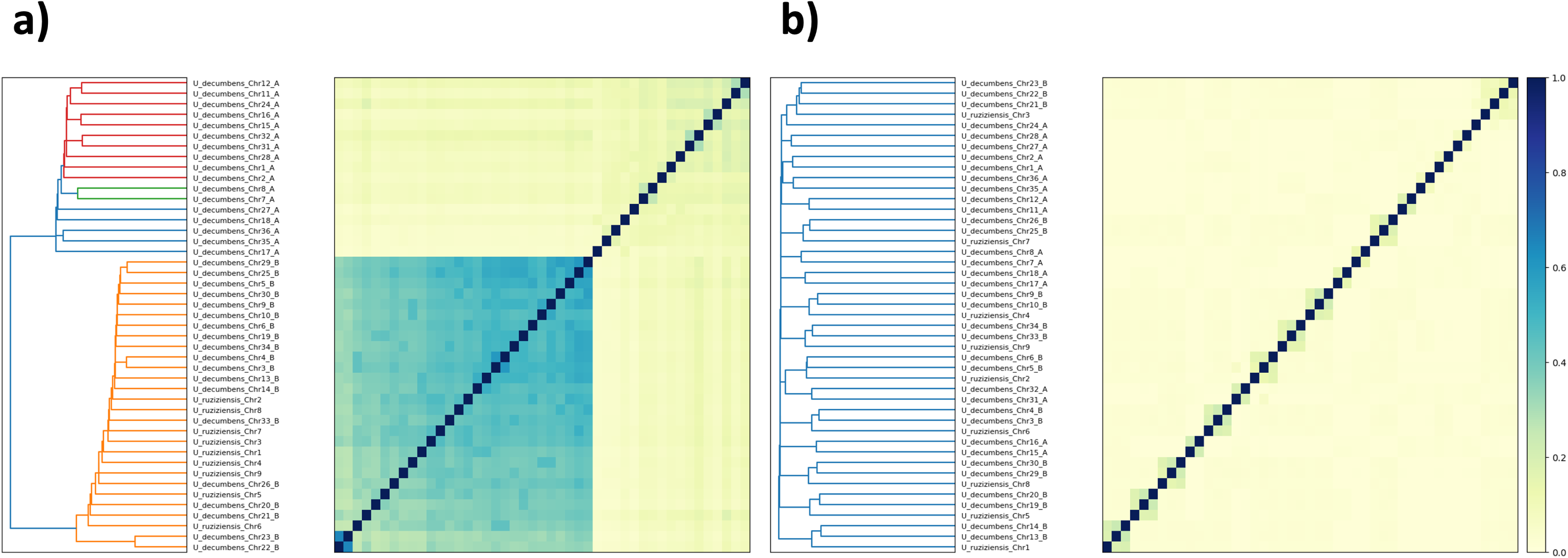
(a) Kmer frequency (K = 81) clustered *U. decumbens* by subgenome (labelled as A and B) and placed all *U. ruziziensis* chromosomes with one of the subgenomes (b) However chromosomes did not cluster by subgenome when using Kmer composition (K=81) instead they evidence preferential pairing restricted within subgenomes, expect in chr. 21 due to a chromosome exchange (also observed by coverage analysis). Branch and heatmap colour represent sequence similarity based on Kmer signatures.

When we grouped the chromosomes from the *U. decumbens* and *U. ruziziensis* genomes together (Figure 4), we noticed that all the *U. ruziziensis* chromosomes clustered with half of the chromosomes that also displayed higher similarity to *U. ruziziensis* in the dotplot. This provides further evidence supporting the relationship of one of the subgenomes in the allotetraploid *U. decumbens* with the diploid *U. ruziziensis*, while the other subgenome has a different origin.

To infer the ancestry of each chromosome we independently aligned WGS short-reads from diploid *U. ruziziensis*, diploid *U. decumbens* and diploid *U. brizantha* to the new assembly: *U. ruziziensis* and *U. decumbens* aligned to the same 18 chromosomes (Figures 5 and S4), while *U. brizantha* reads aligned to the other 18 chromosomes (Figure 5). We concluded that half the chromosomes’ ancestry was from *U. brizantha,* while the other half was from either *U. ruziziensis,* diploid *U. decumbens* or their common ancestor. We could not distinguish between these two species, as there was no difference between where reads from diploid *U. decumbens* and *U. ruziziensis* aligned (Figure S2).

**Figure 5:**
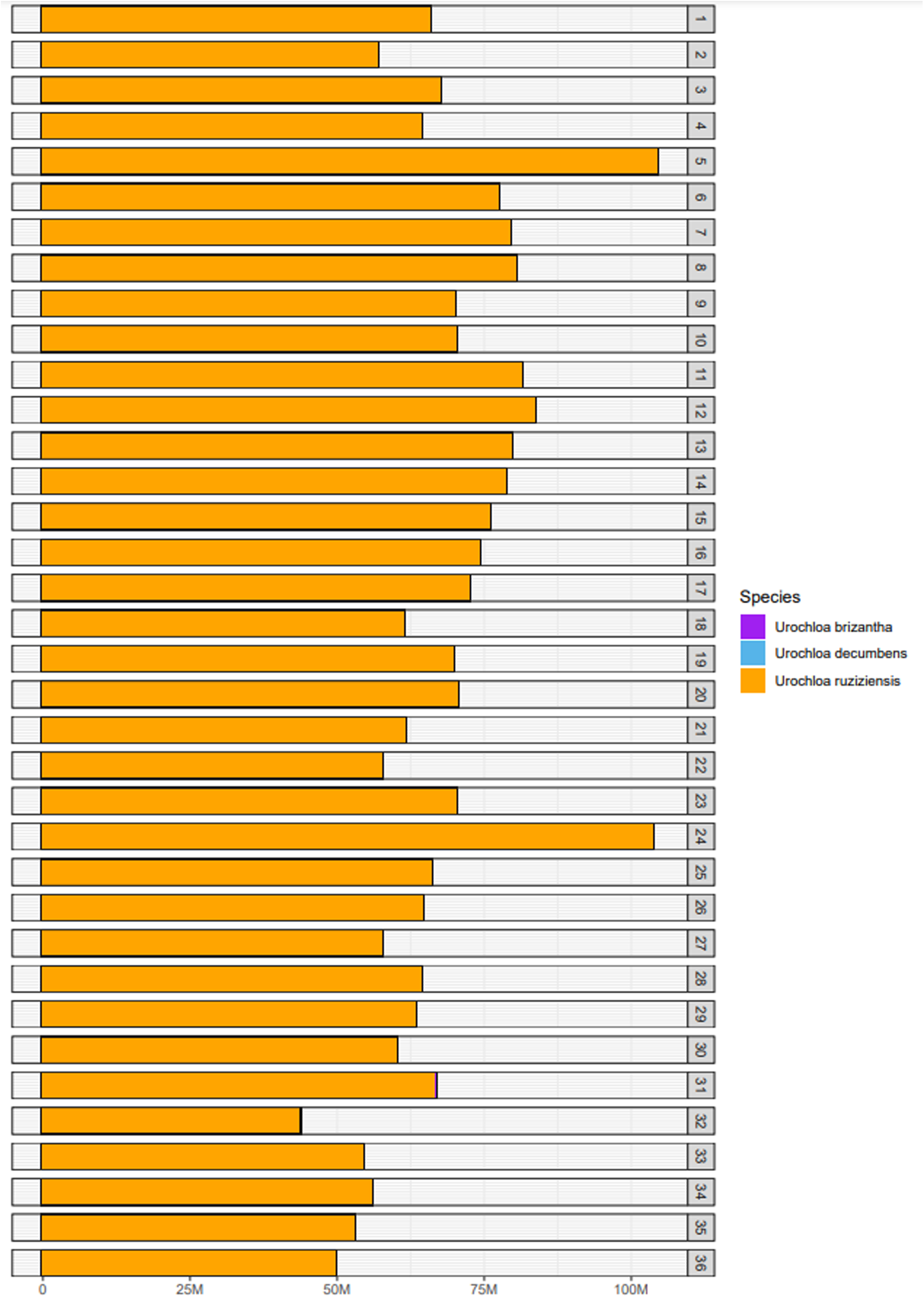
Coverage (read depth) following the alignment from three diploid Urochloa species to the *U. decumbens* genome. Half of the chromosomes were contributed from *U. brizantha* (purple) or a close ancestor, and the other half from diploid *U. decumbens* (blue) or *U. ruziziensis* (orange) or a close ancestor. Evidence for homoeologous exchange (chr. 21) and translocations (chrs. 5 and 32) can also be observed.

Previous phylogenetic and ancestry analyses have shown diploid *U. decumbens* more closely related to diploid *U. ruziziensis* than polyploid *U. decumbens* (Higgins et al. 2022). Another study using Fluorescence In Situ Hybridization (*FISH*) proposed an ancestry of nine chromosomes from *U. brizantha*, nine chromosomes from *U. decumbens* and eighteen chromosomes from *U. ruziziensis* (Tomaszewska et al. 2023). This result likely reflects the difficulty of designing markers that do not cross-hybridise among these highly related species in the *Urochloa* species complex.

On the other hand, it had been suggested that tetraploid *U. decumbens* was a segmental allopolyploid based on genetic mapping (Worthington et al. 2016). However, we did not observe evidence of frequent pairing across subgenomes and homoeologous exchanges in the contact matrix (Figure 2) or coverage plot (Figure 5) to justify its classification as segmental allotetraploid. The only evidence of exchange between homoeologous pairs is chromosome 21 (Table S2), as observed in Kmer composition analysis (Figure 4b) and coverage analysis (Figure 5). Chromosome 21 corresponds to chromosome 8 in *Setaria italica*. This is the same base chromosome (chr. 8) detected in the genetic maps in Worthington *et al*. (2016) that we think led to *U. decumbens’* “historical” classification as segmental allotetraploid. However, our results support this is exclusive to cv. Basilisk and not the whole species.

Finally, we also observed one translocation between chromosomes 5 and 32, where the beginning of chromosome 32 (*U. brizantha* ancestry) had been translocated to the end of chromosome 5 (*U. decumbens/ruziziensis* ancestry) in the cultivar Basilisk, which was collected from the wild and consequently reflects the variation the be expected in a wild apomictic lineage in these complex (Figure 6).

**Figure 6:**
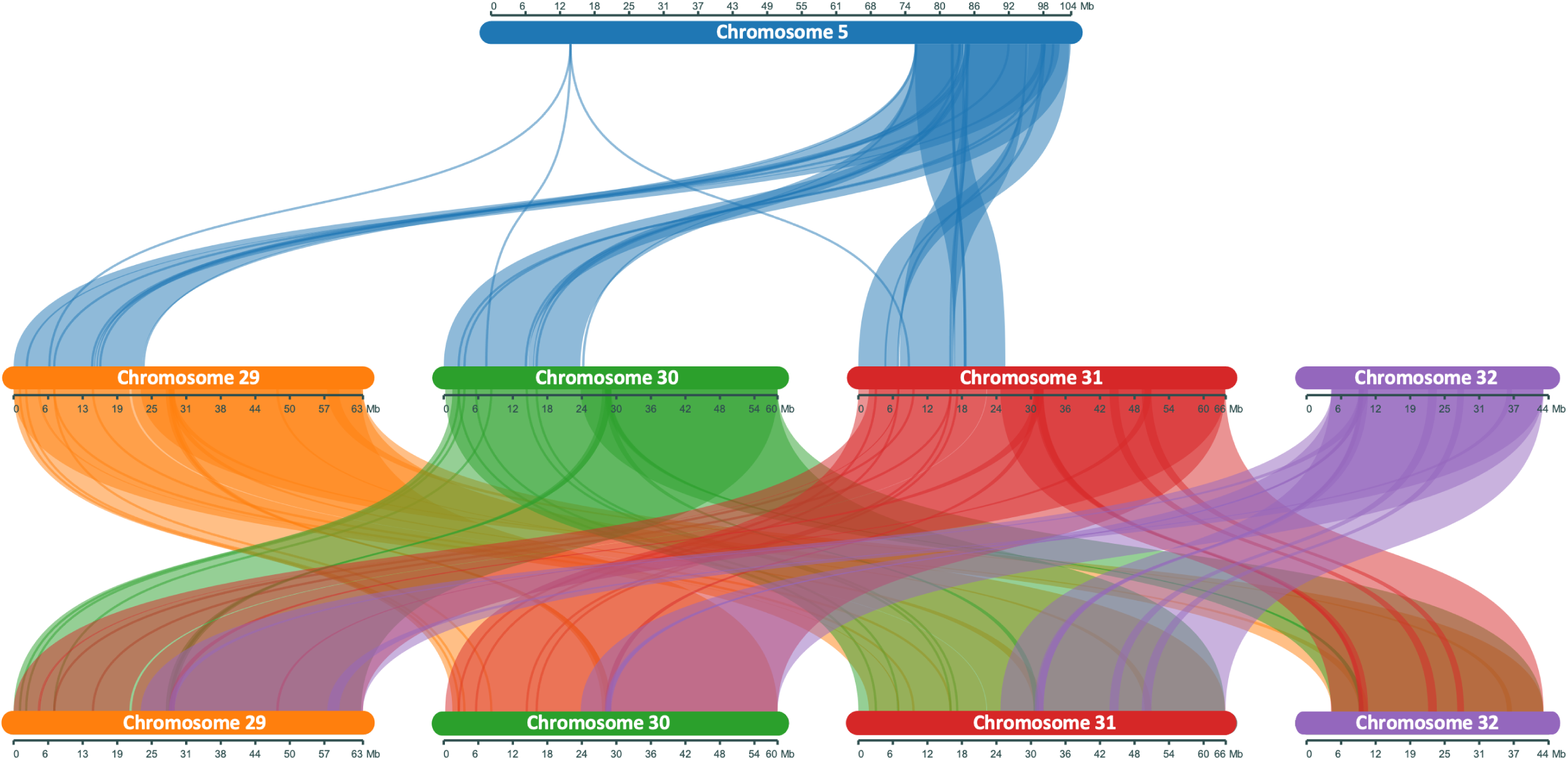
Synteny between chromosomes evidences a translocation between the 5-end of chromosome 32 and the 3-end of chromosome 5. Synteny is conserved between chromosomes 29, 30 and 31, but not 32.

**Figure 7:**
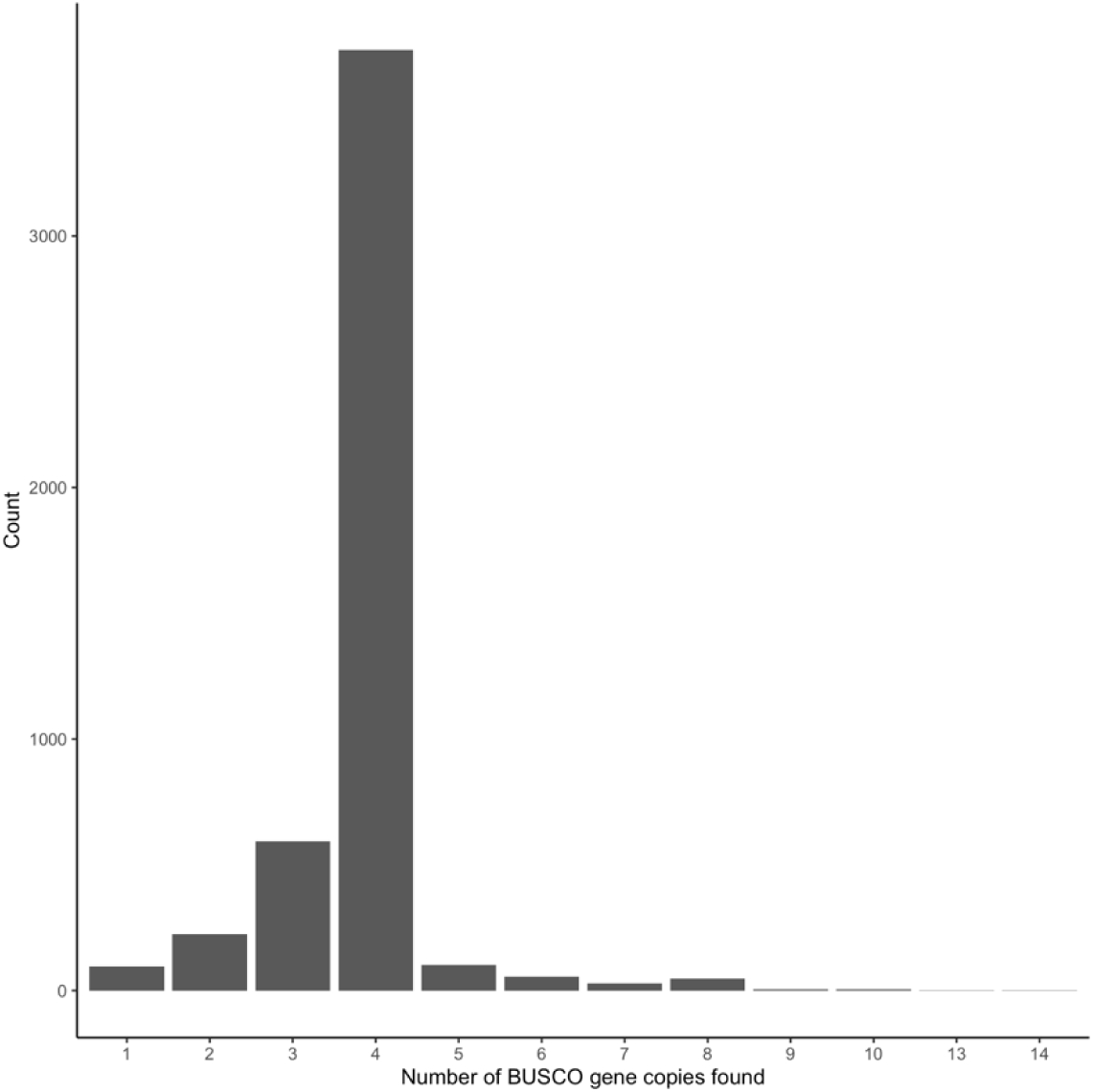
Histogram of the number of times BUSCO single-copy markers aligned in the assembly (Chromosomes only). Most single-copy markers were found four times, once per haplotype, as expected in a heterozygous tetraploid.

Finally, EDTA predicted 2,549,471 interspersed repeats covering 68.35% of the genome (Table 2). The distribution of LTRs across the chromosomes also supported the division of the assembly into its distinct subgenomes, with homologous chromosomes sharing a more similar pattern of repeats than homoeologous chromosomes (Figure 8), except in chromosomes 21-24, where we previously identified a homoeologous exchange in chromosome 21.

**Figure 8:**
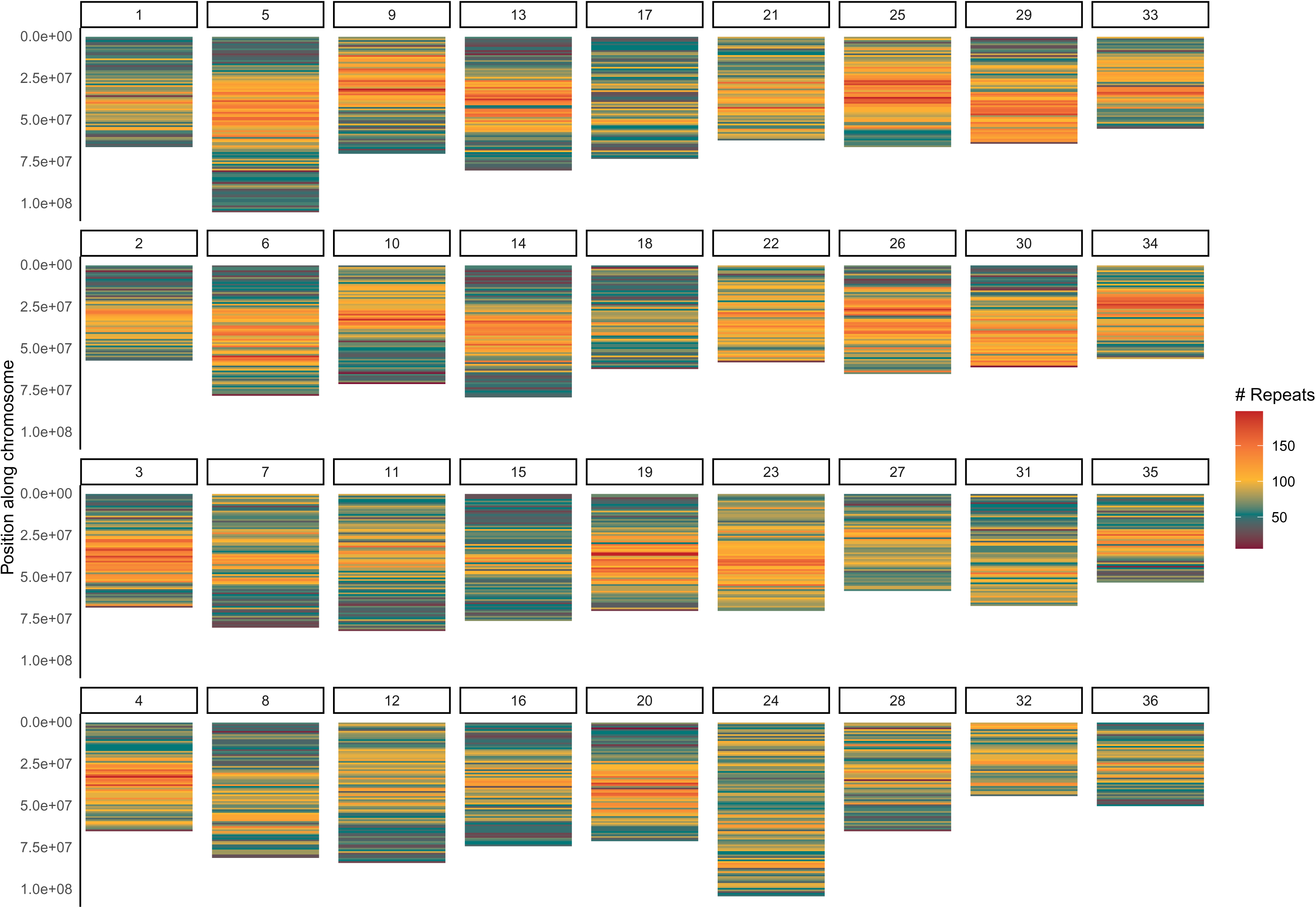
Density of LTRs across the 36 assembled chromosomes evidenced repeat patterns were similar in chromosomes from the same subgenome, except in chromosomes 21-24 due to a homoeologous exchange.

**Table 2:** Summary of repeats regions found in the genome assembly.

| **Class** | **Count** | **Basepairs** | **Percentage/%** |
| --- | --- | --- | --- |
| **LTR** |  |  |  |
| *Copia* | 250,646 | 247,799,132 | 8.61 |
| *Gypsy* | 461,898 | 677,745,937 | 23.54 |
| *unknown* | 405,127 | 424,732,747 | 14.75 |
| **TIR** |  |  |  |
| *CACTA* | 210,015 | 102,760,281 | 3.57 |
| *Mutator* | 181,651 | 70,232,389 | 2.44 |
| *PIF_harbinger* | 106,529 | 36,236,564 | 1.26 |
| *Tc1_mariner* | 163,519 | 46,738,014 | 1.62 |
| *hAT* | 572,94 | 20,684,037 | 0.72 |
| **nonTIR** |  |  |  |
| *helitron* | 712,792 | 341,135,300 | 11.85 |
| **Total interspersed repeats** | **2,549,471** | **1,968,064,401** | **68.35** |

### Conclusion

In this study, we have produced a haplotype-aware chromosome-level assembly of the heterozygous allotetraploid *Urochloa decumbens* cv. Basilisk, an apomictic genotype, using HiFi Pacbio long-reads and Hi-C reads. These technologies have enabled the assembly of all 36 chromosomes of *U. decumbens* at a contiguity functional to the agronomic and scientific community. We also validated the removal (pruning) of Hi-C links between allelic haplotypes, which were innovatively detected using single-copy BUSCO markers, facilitating the anchoring of this haplotype-aware polyploid genome. Furthermore, this haplotype-aware assembly allowed us to identify the ancestry of each subgenome within *U. decumbens*. We concluded that the allotetraploid *U. decumbens* resulted from the hybridisation of diploids from *U. brizantha* and either *U. ruziziensis* or *U. decumbens*. Furthermore, we did not find supporting evidence for its classification as a segmental allopolyploid but for the nominal preferential pairing within subgenomes in allopolyploids. Finally, we believe only haplotype-aware assemblies accurately capture the allelic diversity in heterozygous species, and they should be the preferred option over collapsed-haplotype assemblies in the future.

## Data availability

Raw reads are deposited in the SRA under accession PRJEB73762. The genome assembly, together with its gene annotation, was deposited in ENA with accession GCA_964030465.2 (https://www.ebi.ac.uk/ena/browser/view/GCA_964030465.2). The scripts used in this study are publicly available in Github (https://github.com/DeVegaGroup/HaplotypeAwareChromosomeLevelAssemblyUrochloaDecumbens)

## Acknowledgments

All the authors contributed and approved this manuscript. The authors would like to acknowledge the support of the Norwich Bioscience Institutes Research Computing team. They would also like to thank the Technical Genomics group at the Earlham Institute, especially Drs. Sacha Lucchini, Kendall Baker, Leah Catchpole, Karim Gharbi, and Chris Watkins for their end-to-end support and roles in Project administration and Supervision. Additionally, the authors express their gratitude to Prof. Neil Hall, Dr. Christine Fosker, and their teams, for their roles in Funding acquisition and Supervision. The authors would like to express their gratitude to the staff members of Dovetail Genomics for their input during this project.

---

Figure S1: Kmer spectra supported the assembly accurately captures the sequence information in the HiFi reads.

Figure S2: Coverage (read depth) following the alignment of reads from (a) *U. decumbens* (blue) and (b) *U. ruziziensis* (orange) evidence both aligned almost in the same target regions.

